# A Darwinian outlook on schistosomiasis elimination

**DOI:** 10.1101/2020.10.28.358523

**Authors:** Frederik Van den Broeck, Joost Vanoverbeeke, Katja Polman, Tine Huyse

**Affiliations:** Department of Microbiology, Immunology and Transplantation, Rega Institute for Medical Research, University of Leuven, Leuven, Belgium; Department of Biomedical Sciences, Institute of Tropical Medicine, Antwerp, Belgium; Department of Biology, University of Leuven, Leuven, Belgium; Department of Biology, Royal Museum for Central Africa, Tervuren, Belgium

**Keywords:** Schistosoma, praziquantel, elimination, parasite refugia, genetic reservoirs

## Abstract

Schistosomiasis is a poverty-related chronic disease that affects over 240 million people across 78 countries worldwide. In order to control the disease, the World Health Organization (WHO) recommends the drug praziquantel against all forms of schistosomiasis. Mass Drug Administration (MDA) programs with praziquantel are successful on the short-term as they reduce the prevalence and infection intensity after treatment, and thus instantly relieve the patient from the burden of its disease. However, epidemiological and genetic studies suggest that current school-based interventions may have little or no long-term impact on parasite transmission. Here, we adopt a Darwinian approach to understand the impact of MDA on the neutral evolution of *Schistosoma* parasites and assess its potential to eliminate schistosomiasis. We develop a finite island model to simulate the impact of repeated treatments on the genetic diversity of schistosome populations locally (within each host, i.e. infrapopulation) and regionally (within all hosts combined, i.e. component population). We show that repeated treatments induced strong and lasting declines in parasite infrapopulation sizes, resulting in concomitant genetic bottlenecks within the treated individuals. However, parasite genetic diversity recovered quickly in a few generations due to re-infection, and there was little or no impact of treatment on the genetic diversity of the component population when treatment coverage was 95% or lower. This was mainly due to parasite infrapopulations of the untreated host individuals that acted as reservoirs of genetic diversity, sustaining the diversity of the component population. Hence, lasting declines in parasite genetic diversity were only observed when coverage of treatment was 100%, resulting in population crashes after a minimum of six treatment rounds. We argue that achieving a full coverage of treatment is highly challenging for most endemic regions in sub-Saharan Africa, and conclude that MDA alone has little potential to achieve elimination within a conceivable time frame. Our results raise skepticism about the current WHO goals of elimination of schistosomiasis by 2025.

## INTRODUCTION

Schistosomiasis is a major public health problem in a large part of the world, caused by parasitic trematodes of the genus *Schistosoma*. Over 240 million people are infected across 78 countries worldwide, of which more than 90% of the cases occur in sub-Saharan Africa where close to 300,000 people die annually from the disease [1,2]. This poverty-related chronic disease affects the physical and mental health of children [3,4] and accounts for about 70 million disability-adjusted life-years lost per year [5], which is more than that of malaria or tuberculosis and almost the same as that of HIV/AIDS [6,7].

Since the discovery of the drug praziquantel (PZQ) in the 1970’s, the global strategy of schistosomiasis control shifted from transmission control with molluscicides to morbidity control with PZQ [8]. Over the past two decades, the World Health Organization (WHO) promoted preventive chemotherapy through mass drug administration (MDA) as a strategic approach of targeted and periodic treatment of populations that are infected, or at risk of infection, in order to reduce or prevent schistosomiasis morbidity [9–11]. MDA delivery is usually undertaken based on community distribution of PZQ without individual diagnosis, primarily targeting school-age children who have the highest infection burden and who can be reached efficiently through schools [12].

In 2001, the 54th World Health Assembly set the minimum target of regular administration of chemotherapy to at least 75%, and up to 100%, of all school-age children at risk of morbidity by 2010 [13]. Since then, there has been a growing global effort towards controlling schistosomiasis through MDA with PZQ [14,15], resulting in a decrease in disease prevalence and morbidity in many regions of sub-Saharan Africa [16–20]. However, persistent transmission hotspots of *Schistosoma* species remained [21,22] as the 2010 coverage target was not achieved: only 8.3% of the 239 million people requiring PZQ received preventive chemotherapy [11]. In 2012, the WHO added more ambitious goals of regional elimination of schistosomiasis in the Americas and country-level elimination in Africa by the end of 2020 [11]. The goals of elimination were subject to debate as the NTD community was divided on whether elimination is feasible through MDA alone or whether other measures (such as snail control and improved health education and sanitation) should be integrated [23–28]. In late 2019, a cross-sectional study including multi-year data from nine national schistosomiasis control programs in Africa revealed that the WHO goals of disease control (defined as a <5% prevalence of heavy-intensity infection) were reached in 6/9 countries for *S. mansoni* and 8/9 countries for *S. haematobium* [29]. However, the WHO goals of elimination (defined as a <1% prevalence of heavy-intensity infection) were reached in only 3/9 countries (Burkina Faso, Malawi and Rwanda) for *S. mansoni* and 0/9 countries for *S. haematobium* [29]. While these observations indicate that elimination may be achievable in some areas, in particular areas with low prevalence [29], they also revitalize the ultimate question whether elimination is ever feasible through MDA alone.

Genetic tools and concepts could support disease elimination efforts [30]. By analogy with species conservation genetics that aim to understand population biology to preserve biodiversity and reverse species declines, disease elimination/eradication genetics examines pathogen populations that are deliberately driven to their extinction by human activity [31]. Using genetic diversity as a proxy for effective population size, genetic surveillance could track changes in pathogen demography over the course of MDA treatment programs, identifying populations that are close to extinction. Treatment is expected to substantially decline the number of adult worms (at least locally). Such population bottlenecks may lead to increased endogamy (i.e. mating among related individuals) and low levels of genetic diversity, resulting in the random fixation of (possibly deleterious) alleles through genetic drift (see [32] for theoretical formulations). These population dynamics could decrease parasite fitness, possibly driving the success of mass treatment programs. At least in experimental infections, it has been proven that PZQ treatment decreases *S. mansoni* genetic diversity and increases endogamy [33]. Over the past decade, a total of seven studies quantified the genetic diversity of natural *S. mansoni* populations in sub-Saharan Africa in response to MDA treatment using microsatellites (repeat sequences in DNA that are used to fingerprint individual parasites) [32,34–38] or partial mitochondrial gene sequences [39]. Most of the genetic studies occurred in regions of high endemicity (≧50% infection prevalence among school age children as determined by parasitological methods) (Table 1). Six of the seven studies observed no change in genetic diversity after one to six rounds of PZQ treatment (Table 1), suggesting that schistosome populations were resilient to repeated praziquantel treatments. Only one study reported a decline in parasite genetic diversity following PZQ treatment in two schools of Tanzania [34]. However, factors other than preventive chemotherapy may have caused the decline in these schools because i) the same study also reported a decline in parasite genetic diversity in a control group of children that had not been previously treated as they were not yet of school age during baseline sampling [34], and ii) a follow-up study five years later demonstrated that parasite genetic diversity had recovered and even surpassed baseline estimates despite two to four treatment rounds [37], suggesting that preventive chemotherapy had little impact on fluctuations in parasite genetic variation.

**Table 1.** List of population genetic studies examining the impact of MDA treatment programs in sub-Saharan Africa on the genetic diversity of *Schistosoma mansoni* populations using microsatellites.

| Country | # Schools | Baseline endemicity | Baseline infection intensity | Baseline genetic diversity | Duration | # Treatments | Decline | Ref. |
| --- | --- | --- | --- | --- | --- | --- | --- | --- |
| Tanzania | 2 | H | M-H<sup>s</sup> | H | 6 months | 1 | Yes | [34] |
|  | 2 | H | H | M | 5 years | 2-4 | No | [37] |
| Uganda | 3 | H | M | H | 2 years | 4 | No | [38] |
| Kenya | 4 | H | H | H | 4 years | 4 | No | [36] |
| Senegal | 1 | H | M | M | 2 years | 6 | No | [32] |
| Brazil | 2 | M | M-H<sup>s</sup> | NR | 6 weeks | 1 | No | [35] |

Endemicity is divided into three categories based on infection prevalence by parasitological methods among school age children: low (<10%), moderate (10-50%) and high ( $\geq 50\%$ ). Infection intensity is quantified based on average eggs per gram: low (<100 epg), moderate (100-400 epg) and high (>400). Baseline genetic diversity as estimated by expected heterozygosity per child: low (<0.4); moderate (0.4-0.7); high (>0.7). <sup>s</sup>baseline infection intensity was moderate in one village and high in the other. L = Low, M = moderate, H = high, NR = Not Reported.

As it stands, there is no conclusive genetic evidence for local extinctions nor declines of *S. mansoni* populations in response to MDA treatment programs. As such, it remains unclear whether and under which conditions a schistosome population could be driven to its extinction. Here, we provide a theoretical framework and develop a finite island model to simulate the evolution of schistosome populations locally (within each host) and regionally (within all hosts combined) in response to artificially induced population bottlenecks. The model was defined in such a way that it allows to investigate how parasite evolution is affected by endemicity and the coverage of treatment, providing insights into the conditions under which MDA could possibly lead to population crashes (i.e. elimination).

## METHODS

### Description of the model

The digenean trematode *Schistosoma* spp. has a complex two-host life cycle with an asexual amplification stage in the snail intermediate host, yielding thousands of clonal cercariae that infect the human final host. The cercariae develop into dioecious worms that reproduce sexually within the veins, producing eggs that are expelled into the external environment through urine or feces. Upon contact with water, eggs will hatch into free-swimming miracidia that infect the intermediate snail host. Note that the parasite obligatory cycles through two hosts each generation and that population growth within the human host can only happen due to new infections (Figure 1). Here, we ignore the asexual amplification stage and focus our model on adult parasites in the human hosts, the target of praziquantel treatments. A study on *S. mansoni* infecting wild rat (*Rattus rattus*) showed that the asexual amplification within snails did not play an important role in the total genetic diversity of the adult stages [40]. Models furthermore confirmed that the effect of the clonal amplification phase on adult population genetics is small when there are high levels of parasite gene flow [41,42], which has been reported within most natural schistosome populations [43–46]. Hence, we believe that ignoring the asexual stage in the snail will not affect the outcome of our study when migration is sufficiently large.

**Figure 1.**
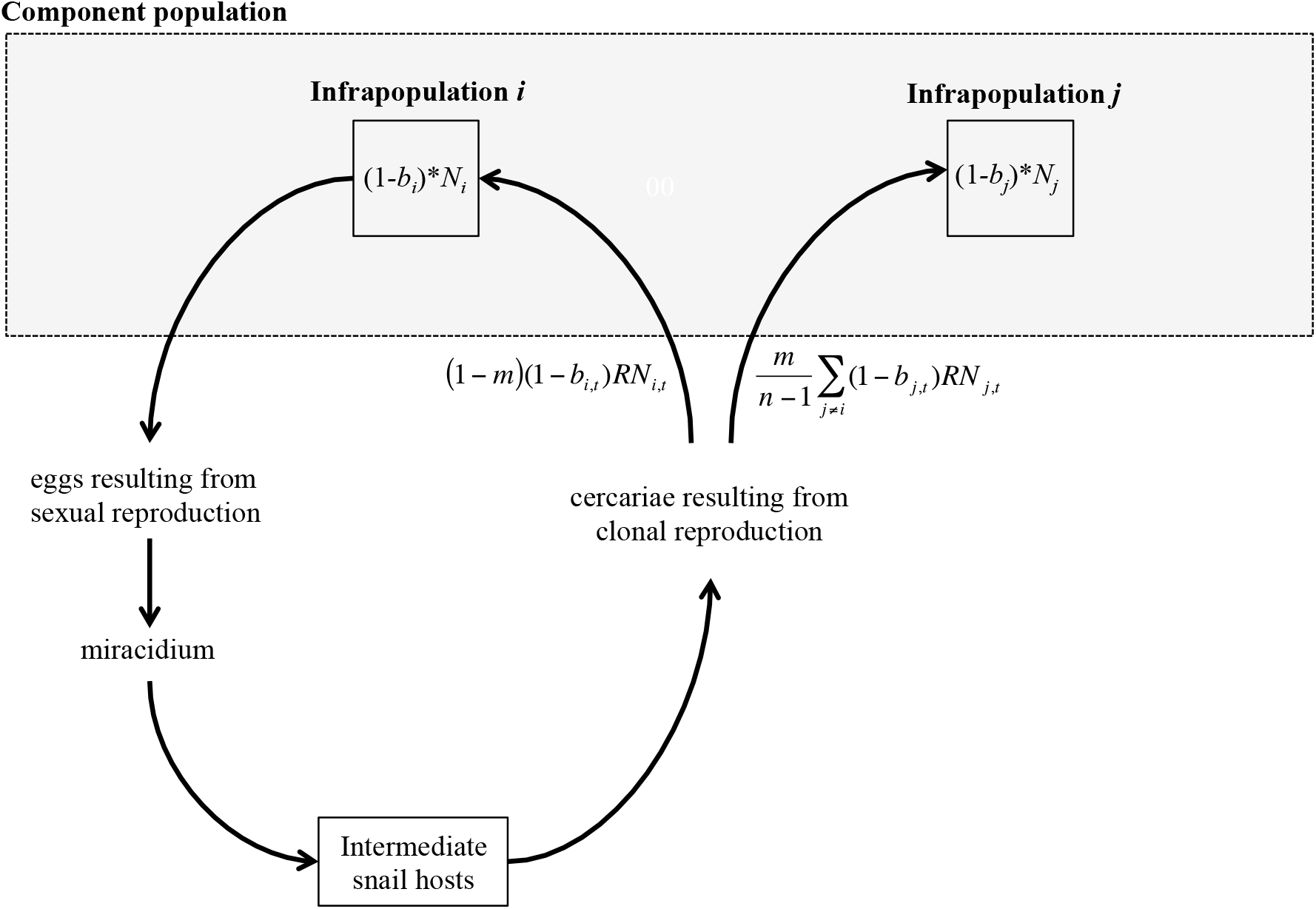
Life cycle of *Schistosoma* spp. with indication of the component population (i.e. all adult worms within one host population) and infrapopulation (i.e. all adult worms within one host individual). Arrows indicate the direction of the life cycle.

We consider a population of 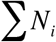 adult worms of the same species within a given host population, i.e. the component population (Figure 1) [47]. The component population represents a given administrative setting (state, region, province, district, village, etc.), but the boundaries of the component populations are not explicitly defined by the model. The component population is subdivided into *n* subpopulations (finite island model), each with a carrying capacity *Kc*. A subpopulation *i* represents a group of *N_i_* randomly mating adult worms of the same species within one individual host, i.e. the infrapopulation (Figure 1) [47]. According to the island model, each reproduction cycle a proportion *m* of the newly produced individuals within a given infrapopulation will randomly infect other infrapopulations, while a proportion 1-*m* infects the infrapopulation of origin. Note that we did not include an explicit spatial structure: gene flow is the same among all infrapopulations and *m* represents the total migration from one infrapopulation to all other infrapopulations. Within each infrapopulation, population growth follows a discrete logistic growth model with per capita infection rate *R*, per capita mortality *b_i_* and carrying capacity *K_c_*.

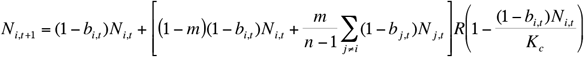

The population size of a schistosome infrapopulation *i* at time *t+1* is thus composed of the individuals that survive treatment and natural death, (1 − *b_i,t_*)*N_i,t_*, plus the individuals that result from new infections by larval stages from parents of the same infrapopulation, (1 − *m*)(1 − *b_i,t_*)*N_i,t_* and from parents from other infrapopulations after migration, 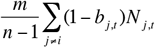.

A given infrapopulation *i* does not decline in size as a result of emigration but only as a result of death, as only larvae migrate and not the adults. *R* represents the per capita number of larvae that successfully infect a human host within the component population, and should not be confused with the basic per capita reproduction number of infective larval stadia. Mortality rate *b_i,t_* comprises both the background mortality (e.g. due to immunity) and the mortality due to drug treatment. The parameter *b_i,t_* can vary over infrapopulations and time to reflect the presence or absence of treatment-related mortality at a given time *t* in infrapopulation *i*. It is defined as a matrix containing mortality rates per infrapopulation *i* (rows) per time unit *t* (columns). We do not explicitly model individuals but rather the set of alleles that represents each infrapopulation. For an infrapopulation of *N* individuals we thus only keep track of the allelic identity of 2*N* alleles without taking into account the genotypic structure of the population. As a consequence, we assume that our infrapopulations are in Hardy-Weinberg and Linkage Equilibrium at all times.

### Implementation of the model

Simulations were initialized by generating alleles in an island model at equilibrium using the function *sim.genot* in the R package ‘hierfstat’ [48]. This function generates alleles following the K-allele mutation model (KAM), and will therefore yield a number of alleles that is consistent for microsatellites. A total of 20 genetic markers were simulated, each mutating to a maximum of 10 possible allelic states. The mutation rate was set to 10e-4, which is the estimated mutation rate for microsatellites in schistosomes [49]. The proportion of migration among islands (i.e. infrapopulations) was set to *m* as used in our subsequent simulations. A total of 100 islands were simulated, a matrix large enough not to result in an artificial loss of regional genetic diversity. The island population size equaled the carrying capacity *Kc*.

The model was initialized three times using different starting values for *Kc* (4000, 500 and 150) to mimic a region of high (mean worm burden *M* = 4288), moderate (*M* = 561) and low (*M* = 147) endemicity, respectively [50]. In other words, we assume that host individuals are infected by a maximum of 150 worms in low endemicity regions, 500 worms in medium endemicity regions and 4000 worms in high endemicity regions. These numbers were chosen based on a predicted distribution of *S. mansoni* worm burden among individuals from three different endemic communities in Burundi and the Democratic Republic of Congo (Figure 1 in [50]).

Once initialized, during each timestep (generation), a number *D* of dying individuals was chosen in each infrapopulation based on a random binomial distribution with *b_i_* probability of mortality. Subsequently, 2*D* alleles were randomly removed from the infrapopulation. After mortality, the number of successful new infections within each infrapopulation was calculated. For each new infection, the infrapopulation of origin was drawn from a random multinomial distribution with probabilities based on *N_i_* and dispersal probabilities, and two alleles were randomly drawn from that infrapopulation based on the current allele distributions.

Drug treatment was implemented by changing the parameter *b_i,t_*, which is defined by a two-dimensional matrix with columns representing populations and rows representing time-steps (generations). Infrapopulations that were not treated experienced the same natural mortality *b_natural_* of 0.04, meaning that 4% of each untreated infrapopulation at each time-step will die due to natural death. This number was chosen because in the absence of infection, a natural mortality of 0.04 would eliminate an infrapopulation within 25 generations, the average lifetime of a schistosome worm (the average lifetime of a schistosome worm is 4-6 years [51], which is equal to 20-30 generations (average 25 generations) assuming an average generation time of 71 days [52]).

Infrapopulations that were treated experienced a mortality *b_PZQ_* of 0.95 assuming that drug treatment would kill 95% of all worms within the human host. In these infrapopulations, the mortality due to natural death was considered negligible in comparison to the mortality due to PZQ. The mortality *b_PZQ_* was not set to 100% because there are many factors that lower PZQ effectiveness, such as i) PZQ is not effective against immature worms present within the host, ii) PZQ is less effective in lightly infected patients that show less robust immune responses [53], and iii) there is variability in PZQ susceptibility among schistosome strains (at least in *S. mansoni*) [54]. Indeed, parentage analyses based on microsatellite and genomic data suggested that adult worms survive treatment [38,55]. The coverage of treatment (i.e. proportion of people that were treated) was implemented by defining *b_PZQ_* for 50%, 90%, 95%, 98% or 100% of the infrapopulations. The frequency of treatment was implemented by defining *b_PZQ_* for a given (set of) infrapopulation(s) once every five generations (assuming an average generation time of 71 days [52]) to mimic the annual treatment administered during MDA programs.

Gene flow *m* was set to 0.99 (i.e. *m* = 1-1/n) to mimic the scenario of a panmictic population, i.e. every host has an equal chance to become infected by parasites from any other host within the administrative setting. This means that our model simulates a group of hosts that share the same gene pool(s), which is a realistic assumption since most studies found low levels of genetic differentiation between parasite infrapopulations from the same household (e.g. [56]) or village (e.g. [57]). Note that the component population is assumed to be completely isolated, i.e. there is no incoming gene flow from other (neighboring) populations.

The parameter *R* was set to 0.5. This value was chosen based on the lifetime reproduction number *R*_0_ that was estimated to be 1-5 for schistosomes [58,59]. For schistosomes, *R*_0_ can be interpreted as the number of mated female schistosomes that were produced by one mated female schistosome during its lifetime. With an average lifetime of *Schistosoma* around 4-6 years [51], reproductive output per generation *R* should be between 0.22-0.9.

Simulations were run for 150 generations. Population genetic statistics were calculated using the allele frequencies after mortality, but prior to new infections. As a proxy for genetic diversity, the number of alleles were estimated at both the infrapopulation level (average number of alleles per locus and per infrapopulation) and the component population level (average number of alleles per locus across all infrapopulations).

## RESULTS

A total of 15 simulations were performed, exploring different scenarios of treatment coverage (50%, 90%, 95%, 98% or 100%) and endemicity (*Kc* = 150, 500 or 4000). At the onset of the simulations (i.e. first generation), infrapopulation sizes averaged 144 worms per host for low endemicity regions, 485 worms per host for medium endemicity regions and 3880 worms per host for high endemicity regions. The median number of alleles ranged between 9.6 and 10 alleles per infrapopulation.

Treatment induced strong declines in infrapopulation sizes (Figure 2A). After the first round of treatment and before re-infection (i.e. fifth generation), the number of worms per treated host dropped from ~144 to ~7 worms in low endemicity regions, from ~485 to ~24 worms in medium endemicity regions and from ~3880 to ~193 worms in high endemicity regions. Parasite infrapopulations then recovered in subsequent generations due to re-infection before the second treatment induced another population bottleneck (Supp. Fig. 1). This cyclic pattern of treatment-induced bottlenecks followed by population recovery was observed during all subsequent generations when coverage was lower than 100% (Supp. Fig. 1). Population sizes of the treated hosts never recovered to baseline levels before the next treatment-induced bottleneck, not even when coverage was only 50% (Figure 2A). For instance, in low endemicity regions and assuming a coverage of 95%, the number of worms would recover to an average 21 worms per host before the next treatment (Figure 2A). Only when coverage was 100%, populations crashed after 30 (six treatments), 35 (seven treatments) and 45 (nine treatments) generations in low, medium and high endemicity regions, respectively (Figure 2A, Supp. Fig. 1). In addition, when coverage was 98%, populations crashed after 90 (18 treatments) and 195 (39 treatments) generations in low and medium endemicity regions, respectively. Populations continued to survive under all other scenarios.

**Figure 2.**
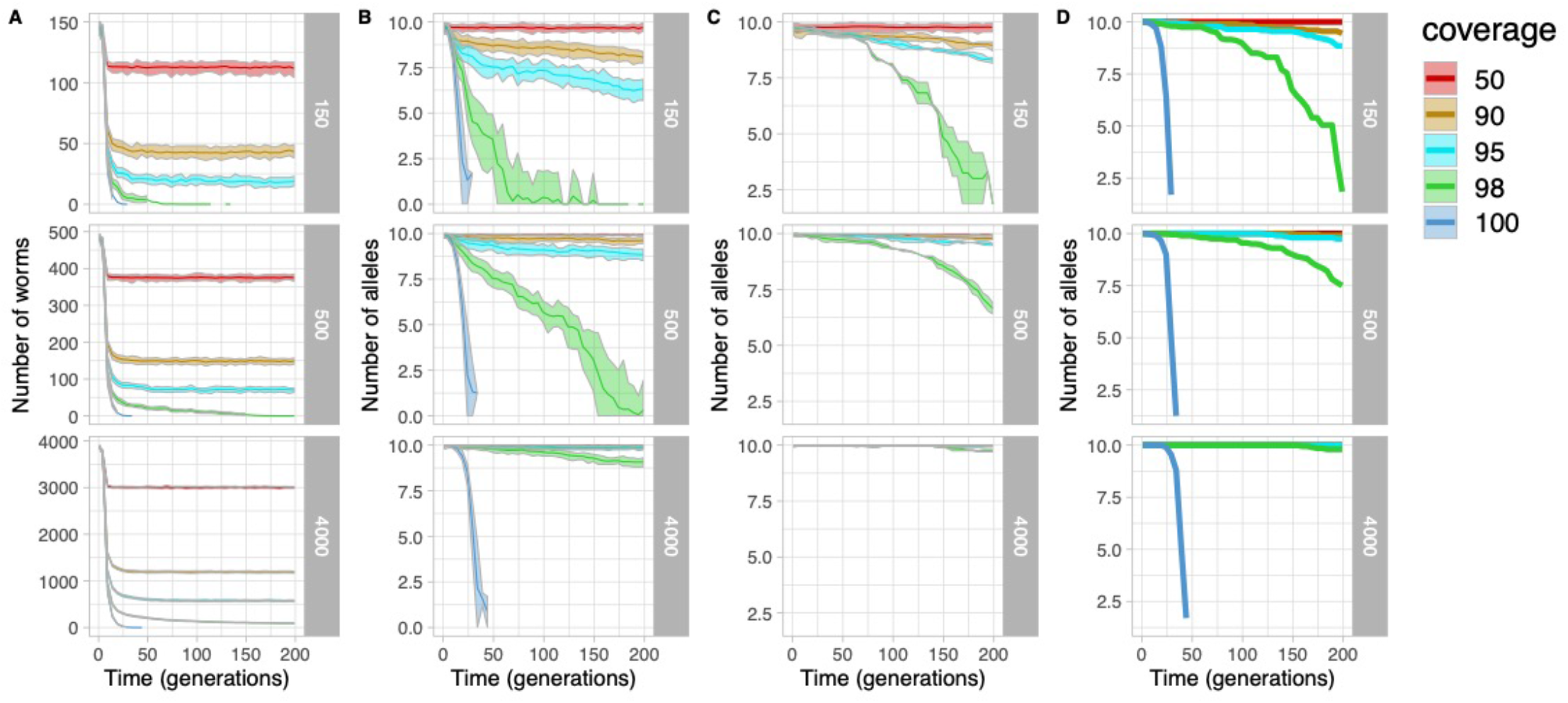
Time-series plots of the number of worms (A) and number of alleles (B-D) in response to repeated treatments every five generations with different treatment coverages (50%-100%) in low (*K_c_*=150), medium (*K_c_*=500) and high (*K_c_*=4000) endemicity regions. For visibility reasons, only numbers as calculated before each treatment were plotted (i.e. generations 1, 2, 3, 4, 9, 14, 19, 24, etc.); complete time-series plots can be found in Supplementary Figures 1-3. (A) Number of worms per infrapopulation. (B) Number of alleles per infrapopulation of treated host individuals. (C) Number of alleles per infrapopulation of untreated host individuals. (D) Number of alleles of the component population. For plots A-C, lines give averages across all infrapopulations, and outer bounds represent the minimum and maximum.

The strong fluctuations in infrapopulation sizes generally resulted in concomitant fluctuations in genetic diversity (Supp. Fig. 2), but the strength of the genetic bottleneck and recovery depended on the endemicity and the coverage of treatment. After the first round of treatment and before reinfection (i.e. fifth generation), the average number of alleles per treated host had decreased from an average 9.69 to 6 alleles in low endemicity regions, from an average 9.95 to 8.43 alleles in medium endemicity regions and from an average 9.99 to 9.77 alleles in high endemicity regions (Table 2). Hence, despite the strong decline in the number of worms per treated host after the first round of treatment, there was not a concomitant decline in genetic diversity in medium and high endemicity regions. In addition, there was virtually no impact of repeated treatments on infrapopulation genetic diversity of treated individuals in high endemicity regions when coverage was 95% or lower (Figure 2B). After 199 generations (just before the 40^th^ treatment), the average number of alleles per host in high endemicity regions was similar to baseline levels when coverage was lower than 100% (Table 2). This was also true for medium endemicity regions when coverage was 90% or lower, and low endemicity regions when coverage was 50% (Table 2). Infrapopulation genetic diversity of the untreated hosts decreased in medium endemicity regions when coverage was 98%, and in low endemicity regions when coverage was at least 90% (Figure 2C). This was not observed in high endemicity regions, where treatment had virtually no impact on parasite genetic diversity in untreated individuals when coverage was lower than 100% (Figure 2C). At the component population level, there was virtually no impact of repeated treatments on the number of alleles per locus in high endemicity regions when coverage was 98% or lower, in medium endemicity regions when coverage was 95% or lower, and in low endemicity regions when coverage was 90% or lower (Figure 2D; Table 2; Supp. Fig. 3). Genetic diversity of the component population was strongly associated with the average infrapopulation genetic diversity of the untreated individuals (Figure 3). Assuming a coverage of 98%, the cross-correlation between the genetic diversity of the component population and the infrapopulations of the untreated individuals was close to one for a time lag of zero, suggesting that at all time points there was a perfect correlation between both variables (Figure 3).

**Table 2.** Average number of alleles per infrapopulation and total number of alleles in the component population, as calculated before and after the first/last treatment. The first treatment occurred at the fifth generation, while the last treatment occurred at the 200^th^ generation. Total loss of alleles due to a population crashes are indicated in gray.

| Carrying capacity | coverage | infrapopulation |  |  |  | component population |  |  |  |
| --- | --- | --- | --- | --- | --- | --- | --- | --- | --- |
|  |  | Before first | After first | Before last | After last | Before first | After first | Before last | After last |
| 4000 | 50 | 10 | 9,84 | 10 | 9,82 | 10 | 10 | 10 | 10 |
| 4000 | 90 | 9,97 | 9,61 | 9,9 | 9,3 | 10 | 10 | 10 | 10 |
| 4000 | 95 | 9,97 | 9,79 | 9,87 | 9,01 | 10 | 10 | 10 | 10 |
| 4000 | 98 | 9,99 | 9,83 | 9,08 | 5,76 | 10 | 10 | 9,8 | 9,8 |
| 4000 | 100 | 10 | 9,74 | 0 | 0 | 10 | 10 | 0 | 0 |
| 500 | 50 | 9,95 | 8,61 | 9,94 | 8,35 | 10 | 10 | 10 | 10 |
| 500 | 90 | 9,97 | 8,42 | 9,62 | 6,7 | 10 | 10 | 9,9 | 9,9 |
| 500 | 95 | 9,98 | 8,52 | 8,85 | 5,13 | 10 | 10 | 9,75 | 9,75 |
| 500 | 98 | 9,94 | 8,3 | 0,31 | 0 | 10 | 10 | 7,5 | 7,5 |
| 500 | 100 | 9,89 | 8,32 | 0 | 0 | 10 | 9,95 | 0 | 0 |
| 150 | 50 | 9,78 | 6,27 | 9,7 | 5,8 | 10 | 10 | 10 | 10 |
| 150 | 90 | 9,62 | 6,09 | 8,06 | 3,42 | 10 | 10 | 9,45 | 9,4 |
| 150 | 95 | 9,68 | 5,96 | 6,34 | 1,91 | 10 | 10 | 8,85 | 8,85 |
| 150 | 98 | 9,74 | 6,13 | 0 | 0 | 10 | 10 | 1,85 | 1,85 |
| 150 | 100 | 9,63 | 5,97 | 0 | 0 | 10 | 9,9 | 0 | 0 |

**Figure 3.**
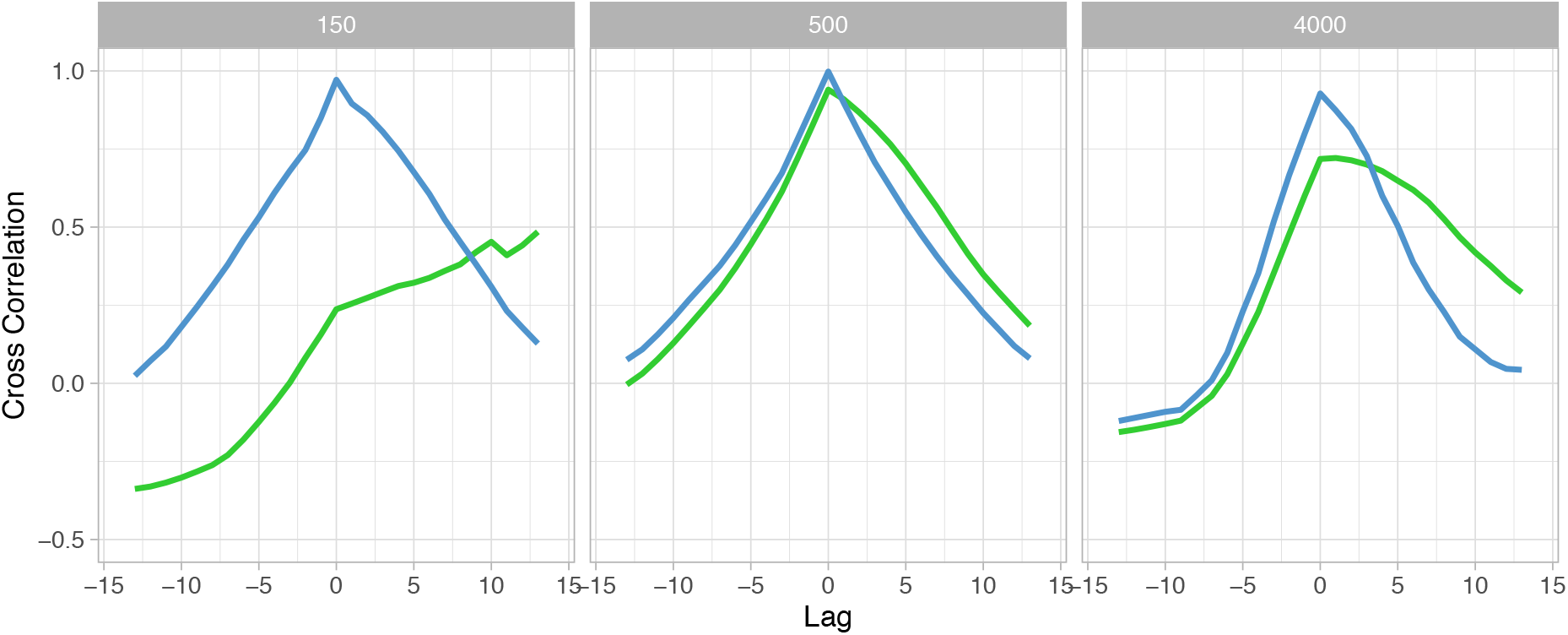
Cross-correlation plots comparing time-series data of the number of alleles found in the component population versus the infrapopulation of the untreated hosts (blue) or the infrapopulation of the treated hosts (green), assuming a treatment coverage of 98%. Only allele numbers as calculated before each treatment were used for estimating correlation statistics (i.e. generations 1, 2, 3, 4, 9, 14, 19, 24, etc.).

## DISCUSSION

We have tracked the neutral evolution of schistosome parasites in response to artificially induced population bottlenecks, mimicking the long-term impact of praziquantel treatment during the course of MDA programs.

Our results revealed that repeated treatments reduced the number of worms per host, which is consistent with epidemiological observations of a decrease in worm burden as a result of current control practices [16–20]. Population bottlenecks resulted in concomitant declines in parasite genetic diversity immediately after treatment. However, genetic diversity recovered quickly in a few generations due to re-infection, in particular in medium/high endemicity regions and when coverage was 95% or lower. Under these conditions, regional genetic diversity did not decline after a total of 40 treatments. These results are consistent with observations of sustained levels of genetic diversity during field-based genetic studies, which targeted school-age children (presumably achieving a coverage lower than 95%) in medium/high endemicity regions (Table 1). Similar to our observations, Gower and colleagues found a significant reduction in adult worm load [37] and Faust and colleagues found a decrease in genetic diversity shortly after treatment [38], while both studies found no decrease in diversity over the entire period of the study. Altogether, these results clearly suggest that treatment has only shortterm impacts - important to reduce the adult worm load of the patient - but not sufficient to affect long-term parasite population dynamics.

Our model provides two related explanations for these observations. First, genetic diversity will only decrease over time in medium/high endemicity regions when the coverage of treatment was at least 95%, a number that is rarely achieved during MDA programs. Indeed, MDA programs focus their campaigns mainly on school-age children, leaving pre-school children, adult individuals, lightly infected individuals and asymptomatic carriers untreated. In addition, it is often impossible to reach all school-age children because of the difficult conditions met in sub-Saharan Africa, such as a low school enrollment rate [60]. For instance, a study on soil-transmitted helminthiases in Kenya estimated that school age children harbor as little as 50% of *Ascaris lumbricoides* worms, and that only 40-70% of these children are enrolled at school [61], suggesting that a relatively small portion of the parasite population was exposed to treatment. Second, genetic diversity of the component population is associated with infrapopulation diversity of the untreated hosts. This is a direct consequence of the population being panmictic: almost all genetic variability present within the component population will be found within each individual infrapopulation. Genetic studies assessing the impact of treatment in natural populations found no significant differences in genetic diversity between hosts and/or villages [32,34–38], a pattern typically observed for *Schistosoma* that supports a panmictic parasite population [45,62]. Hence, untreated individuals will not only act as continued reservoirs of transmission [63–65], but also act as reservoirs of regional genetic diversity. Adult hosts in particular may be suitable reservoirs of genetic diversity because their parasite infrapopulations are genetically more diverse compared to those of children [57]. Our model thus suggests that community-wide distribution of praziquantel is essential to induce a long-term impact on parasite population dynamics [66].

The WHO has set elimination as a public health problem (defined as a <1% prevalence of heavy-intensity infection) as a goal for schistosomiasis by 2025. Here, we adopt a similar definition of elimination but define a heavy-infection intensity when the number of worms is equal or larger than 100. Adopting this definition, we found that elimination occurred after one or two treatments when coverage was 100% (Table 3). When coverage was lower than 100%, elimination didn’t occur in high endemicity regions after 40 rounds of treatments. In medium endemicity regions, elimination would happen after 25 rounds of treatment when coverage was 98%, but it would not occur after 40 treatment rounds when coverage was 95% (Table 3). These results suggest it is highly unrealistic to achieve elimination of schistosomiasis through MDA alone. Only when all infected hosts within a given administrative setting received repeated treatments (i.e. 100% coverage), and assuming no immigration of parasites from other administrative settings, elimination will be achieved within a conceivable time frame. However, full coverage is rarely achieved in natural conditions because of parasite refugia, namely the group of parasites that are not reached during a control program [30]. Beside the untreated host individuals (see above), there are other possible refugia such as are larval stages within infested water or intermediate snail hosts, immature worms that escape treatment within human hosts [67], adult worms that survive treatment [38] and alternative reservoir host species such as rodents or baboons that are not reached during MDA [68,69]. In addition, studies have shown a substantial decline of compliance over prolonged chemotherapy campaigns (leaving hosts uncured) [70–72], and of praziquantel efficacy in schools with higher exposure to MDA (leaving worms surviving treatment) [73]. These observations indicate that it is highly challenging to achieve 100% treatment coverage in a given administrative setting. Our model thus suggests that - at least within medium/high endemicity regions - current praziquantel-based control programs do not have the potential to eliminate schistosomiasis. We advocate that integrated control programs including praziquantel treatment, health education and WASH (water, sanitation and hygiene) will have a much higher chance to eliminate schistosomiasis compared to MDA alone [23,28].

**Table 3.**
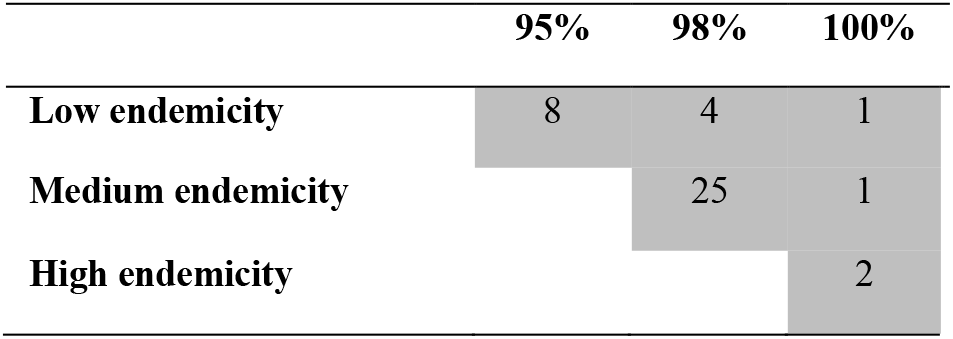
Number of treatments needed to achieve elimination as a public health problem (ELPH) in low, medium and high endemicity regions. Here, elimination was defined as <1% heavy-infection intensity (number of worms is equal to or larger than 100). Numbers are only given for treatment coverages of 95% or higher. ELPH was not recorded after 200 generations and 40 treatments when coverage was lower than 95%. In addition, ELPH was not recorded in high endemicity regions when coverage was 95% and 98%, and in medium endemicity regions when coverage was 95%.

---

**Supplementary Figure 1.**
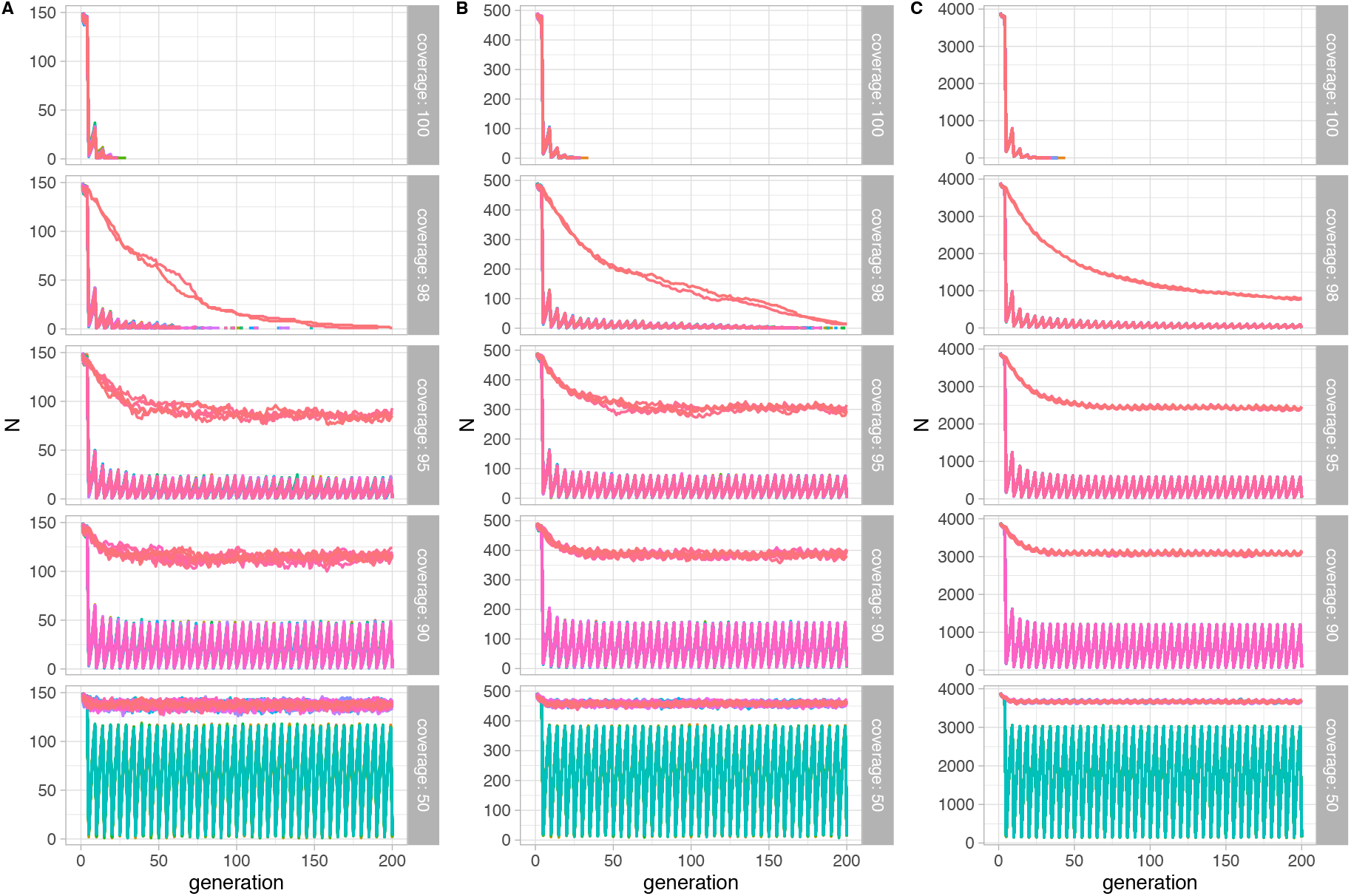
Time-series plots of the number of adult worms per infrapopulation for different treatment coverages (50-100%) in low (**A**), medium **(B**) and high (**C**) endemicity regions.

**Supplementary Figure 2.**
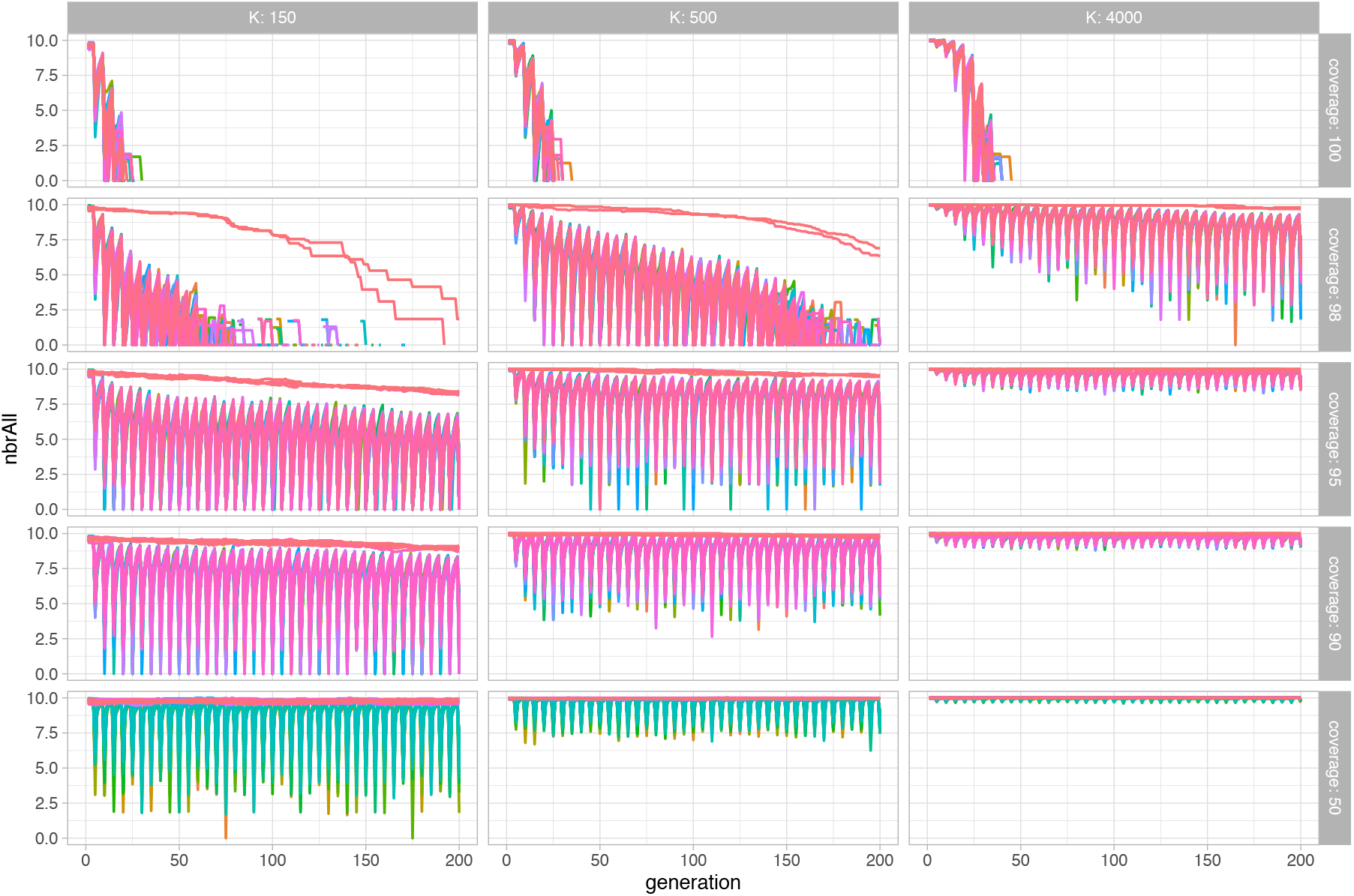
Time-series plots of the number of alleles per infrapopulation for different treatment coverages (50-100%) in low (***K*_c_=150**), medium (***K*_c_=500**) and high (***K*_c_=4000**) endemicity regions.

**Supplementary Figure 3.**
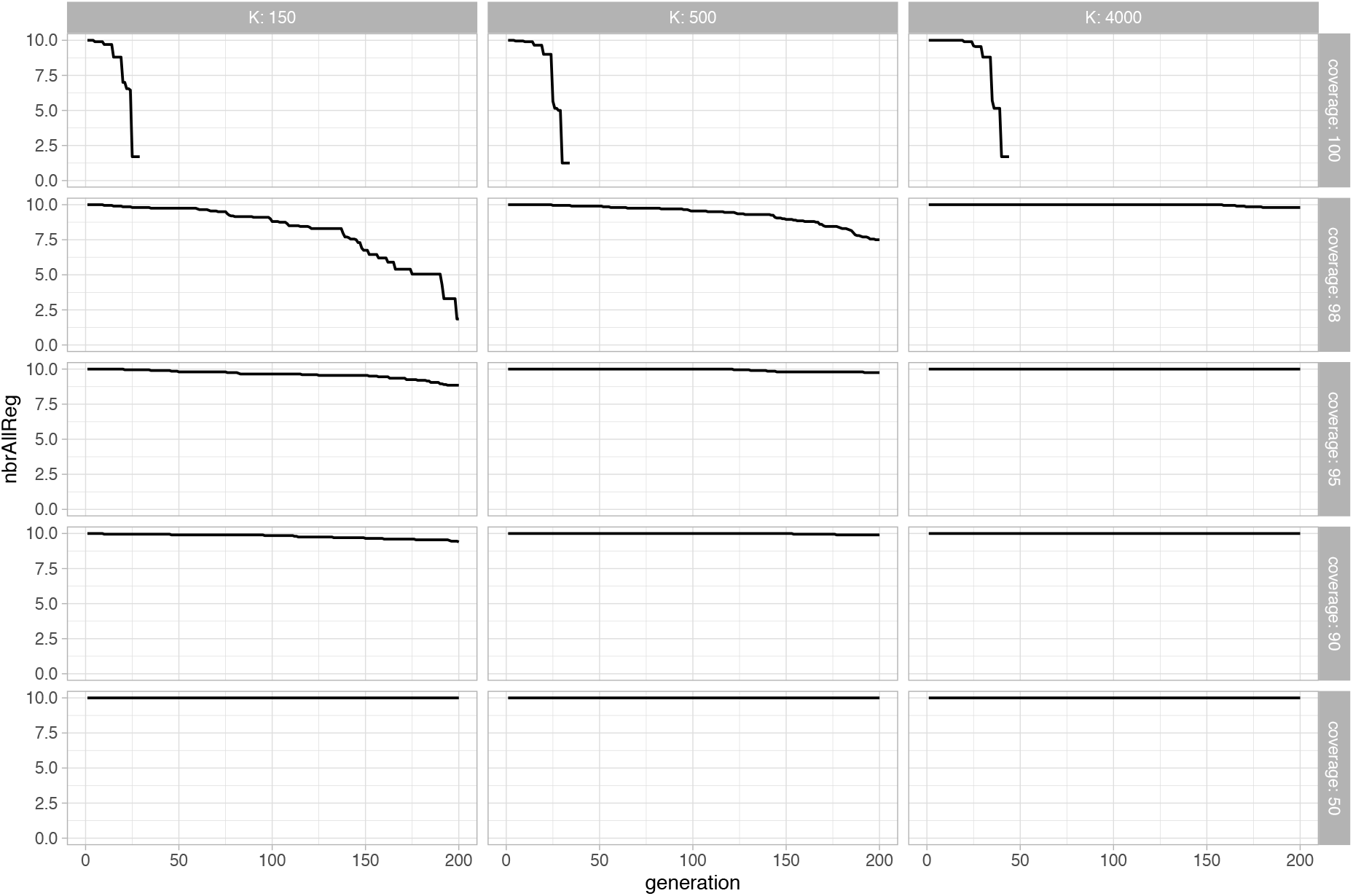
Time-series plots of the number of alleles per component population for different treatment coverages (50-100%) in low (***K*_c_=150**), medium **(*K*_c_=500**) and high (***K*_c_=4000**) endemicity regions.

